# Flight performance, activity and behaviour of breeding pied flycatchers in the wild, revealed with accelerometers and machine learning

**DOI:** 10.1101/2024.03.21.586090

**Authors:** Hui Yu, Shujie Liang, Florian T. Muijres, Jan Severin te Lindert, Henrik J. de Knegt, Anders Hedenström, Koosje P. Lamers, Per Henningsson

## Abstract

Flight behaviours have been extensively studied from different angles such as their kinematics, aerodynamics and more general their migration pattern. Nevertheless, much is still unknown about the daily flight activity of birds, in terms of their performance, behaviour and the potential differences between males and females. The recent development of miniaturized accelerometers allows us a glimpse into the daily life of a songbird. Here, we tagged 26 pied flycatchers (*Ficedula hypoleuca*) with accelerometers and analysed using machine learning approaches their flight performance, activity and behaviour during their chick rearing period. We found that during two hours of foraging chick-rearing pied flycatchers were flying 13.7% of the time. Almost all flights (>99%) were short flights lasting less than 10s. Flight activity changed throughout the day and was highest in the morning and lowest in the early afternoon. Male pied flycatcher had lower wing loading than females, and peak flight accelerations were inversely correlated with wing loading. Despite this, we found no significant differences in flight activity and performance between sexes. This suggests that males possess a higher potential flight performance, which they not fully utilized during foraging flights. Our results thus suggest that male and female pied flycatcher invest equally in parental care, but that this comes at a reduced cost by the male, due to their higher flight performance potential.

## Introduction

Bird flight behaviours have been studied for many years, including detailed examinations of flight kinematics (e.g., Hedenström and Møller, 1997, Krishnan et al., 2022), aerodynamics (e.g., Muijres et al., 2012a, Alerstam et al., 2007) and general aspects such as migration flight strategies (e.g., Mitchell et al., 2015, Jiguet et al., 2019). Despite this, still very little is known about the fundamental aspects of what is required from a bird in terms of flight performance in their day-to-day lives during routine transport flights and foraging flights. In the daily life of songbird, foraging occupies a large portion of the time. To forage, the bird must move through its habitat, which is done predominantly by flying. The amount of food that the bird needs to eat during a day varies through the year (Evans et al., 1994). One of the most demanding periods of the annual cycle may be during breeding and, in particular, during the critical chick feeding phase. This intense period puts great demands on the flight performance of the bird. Despite the crucial importance of flight performance for birds, very little is known about the activity patterns and performance of birds in the wild. This originates from the logistical challenge of studying detailed behaviours in the field, posing a gap in our knowledge about bird behaviour in the wild.

In this study we investigate the flight activity, behaviour and performance of pied flycatchers (*Ficedula hypoleuca*) during chick rearing. Pied flycatcher has an active foraging behaviour, feeding on live insect prey, often captured mid-air (Böhm and Kalko, 2009). This foraging behaviour may involve high performance flights with high demands for agility and endurance, making these birds ideal for studying activity patterns and flight performance. A lot is known about pied flycatcher kinematics and aerodynamics from previous wind tunnel studies (Muijres et al., 2012a, Muijres et al., 2012b, Johansson et al., 2018) which is relevant when attempting to understand and explain their performance and behaviours in the wild.

Furthermore, we also have great opportunities to study this species due to other ongoing projects that involve monitoring the many nest boxes in our study area (e.g., Lamers et al., 2023). Breeding behaviour in this population was therefore closely monitored, including information on hatching date, brood size, identity of individuals and breeding pairs through metal rings. This made it possible to consistently time our measurements in relation to breeding stage, allowing for subsequent comparisons between individuals.

Traditionally, study of insectivorous birds relied on direct human observations (e.g., Alatalo et al., 1982, Alatalo and Lundberg, 1984). However, using new technology, it has been possible to study birds remotely through e.g. radar (Alerstam et al., 2007), radio telemetry (Taylor et al., 2017) and due to recent technological developments using on-board miniature logging devices (e.g., Ropert-Coudert and Wilson, 2005, Stidsholt et al., 2018). Several behavioural and positional parameters can be recorded by loggers attached to animals with little disturbance to their daily activities. Accelerometers, which record accelerations over time, allow the study of animal activities without the limitation of visibility and observer bias (Brown et al., 2013). Accelerometers can be used to record coarse flight activity patterns of birds over a complete annual cycle (Bäckman et al., 2017, Norevik et al., 2019, Macias- Torres et al., 2022). What these accelerometer studies do not show, however, is activity patterns and detailed flight performance at high temporal resolution. Recent application using supervised machine learning methods (i.e., where behaviour types from direct observation of the tracked individual are necessary for model training) had advantages for identification of complex behaviours such as food ingestion in spoonbills (Lok et al., 2023). Therefore, we applied a similar supervised machine learning model on pied flycatchers (Yu et al., 2023) in this study. In addition, dynamic body accelerations (i.e., the acceleration caused by animal body movement) derived from raw accelerometer measurements are widely used as a proxy for energy expenditure (Wilson et al., 2006, Gleiss et al., 2011, Wilson et al., 2019, Sutton et al., 2023). Therefore, we used VeDBA (Vectorial Dynamic Body Acceleration) in this study for flycatcher flights efforts and flight performance comparison. In addition, flight performance of birds was found related to wing morphology (Hedenström and Møller, 1997). Therefore, we quantify the wing morphology and investigate how it relates to flight performance and if there is a difference between males and females.

Furthermore, we compare the investment between males and females for chick care, to study the division of labour between the sexes. Game theoretical analysis show that in monogamous pairs where both parents feed the young, any change in feeding effort by one member of the pair should be countered by a change by the other (Houston and Davies, 1985). With realistic slopes of the reaction curves the resolution may be an evolutionary stable strategy where both parents invest equally much in feeding, whereas the total effort by the couple should increase with increasing clutch size (Houston and Davies, 1985). Finally, we look at how brood size may affect the investment of the parents in terms of flight effort.

## Materials and Methods

### Miniature accelerometers

The accelerometer device (Figure 1) used for this study was designed and developed by the Electronics lab at the Department of Biology, Lund University, Sweden. It is a small device (18×9×2mm, W×L×H) and it weighs 0.7g. The logger contains a LED light, a light sensor (for activation), a micro-electromechanical system (MEMS) accelerometer, a processor, a zinc-air button cell (A10, 100 mAh capacity) and a non-volatile memory. The accelerometer unit was set to record three-axis acceleration vector (**A**=(*a_x_,a_y_,a_z_*) in *g*-force where 1*g* equals 9.81 m/s^2^, with the *x*-axis in the lateral direction, *y*-axis longitudinal, and *z*-axis vertical), at a sampling frequency of 23Hz for the field measurements and 100Hz for the aviary measurements. The measurement range was set to ±8*g* with 8-bit output resolution for each axis, which gives 256 levels and thereby 0.063*g* resolution. The loggers were programmed to include a 30-minute delay from activation until the start of data collection to allow birds to resume activities after being handled. Once sampling had started the device recorded the birds’ accelerations continuously for approximately two hours in the 23Hz configuration and 30 minutes in the 100Hz configuration, limited by memory size which could store approximately 175 000 individual 3-axis recordings.

**Figure 1.**
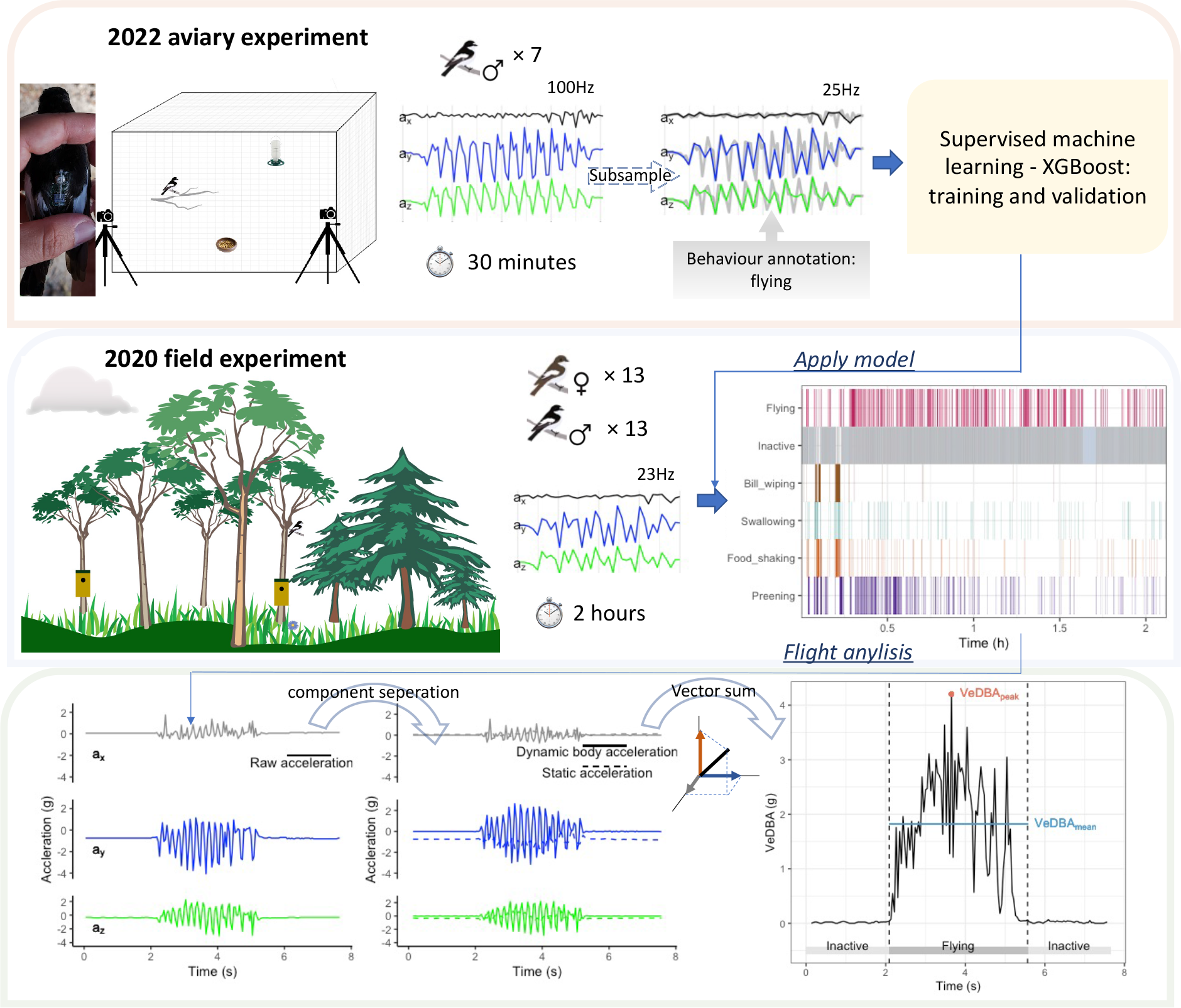
Diagram of experimental setups in two years. In 2022, seven male pied flycatchers undertook aviary experiments. Each of them was tagged with an accelerometer logger. Activities of each bird were recorded by the accelerometer logger as well as two cameras for 30 minutes. Accelerometer data were subsampled from 100 Hz to 25 Hz to be comparable to the 2020 field experiment sampling frequency. Then the accelerometer data were annotated to 6 behaviour types through video records and the annotated dataset was used to train an XGBoost model. In 2020, 13 male and 13 female pied flycatchers were tagged with the same accelerometer loggers as in 2022. Each logger recorded accelerometer data at 23 Hz for around 2 hours duration when the tagged bird moved freely in their natural environment. The XGBoost model from 2022 was used to predict behaviour types of birds tracked in 2020.

### Field measurements

Field measurements were performed in June at Vombs fure, Lund, Sweden, a forest habitat where around 400 monitored nest boxes are available for passerines to breed in (coordinates in decimal degrees: 55.66301, 13.55550) (Figure 1). We successfully recorded activity for 26 pied flycatchers in total, 13 males and 13 females. The birds were caught at their nest boxes using efficient and humane methods, primarily through spring-loaded aluminium trap doors at the entrance hole. Secondarily, if an individual for some reason was reluctant to enter the nest box, we used mist nets put up in front of the nest box. Adult birds were caught when chicks were seven days old to ensure that the chicks could cope with the brief disturbance. It has been shown that the weight of chicks usually reaches an asymptote around day seven (Lundberg and Alatalo, 2010), suggesting that they are from here on less vulnerable to disturbance. The handling of the adult birds for measuring, attaching and removing the loggers took a few minutes and the total time they carried the loggers was typically about five hours. Each individual was measured only once. Hence, the disturbance to these birds was brief in relation to the full breeding period. Brood size and nestling mass were recorded at day 7 and day 12 to record growth rate.

The accelerometers were placed over the synsacrum of the adult flycatchers using a leg-loop harness (Rappole and Tipton, 1991), made from 0.7 mm elastic wire. The battery, albeit very small and lightweight, is the heaviest component of the logger, so to balance the weight of the accelerometer it was mounted with the battery end towards the front. This way the heaviest part of the logger is close to the centre of mass of the bird, which should minimize any detrimental pitch moment and its influence on flight performance.

To record wing morphology of each individual, we took photos of the wings using a ruler as a reference scale in the photo, where one wing of the bird was spread manually. Weight of the bird with logger was recorded. After we had deployed the logger on the bird and measurements were taken, we immediately released the bird close to its nest box.

The logger was retrieved later the same day (n=17) by catching the individual again (7 loggers were retrieved on the following day and 2 were retrieved 2 days after, due to failure to re-capture in the same day). Data were later downloaded after which the logger could be reused. In total we had 12 loggers at our disposal. The average body mass of our flycatchers was 12.5g, so the logger weight was approximately 5% of this. The capture and experimental protocols were approved by Malmö— Lund University Animal Ethics Committee (Permit Nos. 5.8.18-05926/2019 and 5.8.18-05284/2022).

### Aviary measurements

To be able to extract detailed behaviours from the acceleration data we applied machine learning techniques. In order to apply a supervised machine learning method that require annotating behaviour types to raw accelerometer data, we did aviary behaviour observations of seven pied flycatchers in June 2022 (Figure 1). Details of the aviary experiment can be found in Yu et al. (2023).

### Machine learning

The details of aviary-based behaviour annotation, and machine learning model training and validation can also be found in Yu et al. (2023). Importantly here, we removed the category “other”, since it only constituted 1% of pied flycatcher time budget and is not helpful in this study. Therefore, six behaviour categories – flying, inactive (also include behaviours when the birds were perching still, e.g., vigilance or searching for food while perching), food shaking, preening, swallowing and bill wiping - were classified by XGBoost machine learning method. Since there was a mismatch between sampling frequency of aviary-based experiments (100 Hz) and the field study (23 Hz), we used subsamples of aviary-based data by taking every fourth data point (3-axial accelerometer) from the original dataset. Each sample window for behaviour classification contained 16 datapoints, which was roughly 0.7 s duration.

The behaviour classification model trained on aviary-based dataset was then applied on the field data (Figure 1). As we are here primarily interested in flight behaviour, we converted all 0.7 s sample windows classification into flying or non-flying. Next, to reduce misclassification, we changed all single non-flying sample windows between two flying ones into flying. Thus, three consecutive sample windows “flying, non-flying, flying” were changed into “flying, flying, flying”. We then divided the filtered flight sequences into a series of flying and non-flying segments, whereby each segment was given its own start time and duration value.

### Data processing and analysis

#### Quantifying flight activity, effort and performance

Based on the flights identified by the XGBoost machine learning model, we derived several flight attributes (Figure 1). We estimate flight proportion (*R*_flight_) as the total duration of flights divided by duration of the full recoding sequence. Mean flight duration (*T*_flight_) was defined as total duration of flights divided by total number of flights. These metrics were used to quantify the flight activity of the studied pied flycatchers.

The corresponding flight effort and performance were estimated based on the in-flight accelerations, as quantified using the Vector of Dynamic Body Acceleration (VeDBA). The calculation of VeDBA followed Qasem et al. (2012). We converted the raw accelerometer data (**A**) recorded by the loggers into Dynamic Body Acceleration (**DBA**) values by first smoothing each axis to derive the static acceleration (**A**_static_) using a running mean over a time duration of 0.22s (i.e., 5 samples at a 23 Hz sample rate), and then subtracting the static acceleration from the raw data (**DBA** = (**A** – **A**_static_)). Then VeDBA values were calculated as

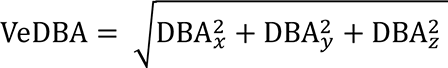

From this, we then calculated the following flight effort and performance parameters per flight segment. We estimated the average flight effort during each flight segment as the mean VeDBA value during that segment (VeDBA_mean_). We estimated the peak flight performance during a flight segment as the maximum value of VeDBA during that segment (VeDBA_peak_).

Based on these mean and peak VeDBA values per flight segment, we estimated the following flight performance metrics per individual bird. The average flight effort per individual was estimated as the mean VeDBA for all flights performed by that individual (VeDBA_mean,bird_).

Mean and maximum flight performance per individual was estimated based on all peak fight accelerations (VeDBA_peak_) estimated for that bird. Mean peak acceleration (VeDBA_peak,mean_) was calculated as the mean of all VeDBA_peak_ values for all flight segments of the individual bird, and quantified the mean flight performance of that bird. The maximum peak acceleration (VeDBA_peak,max_) was equal to the maximum VeDBA_peak_ value among all flight segments of the individual bird, and quantified the maximum flight performance exhibited by that bird.

#### Wing and body morphology

We determined the mass *m* (kg) of all birds by weighing them using a Pesola spring scale, and we characterized the wing morphology using photos of the wings captured at the time of logger attachment. From these photos, we determined single wing area (m^2^) and semi-span (m) using ImageJ (National Institutes of Health, USA), and following the procedure established by (Pennycuick, 1989). We then doubled these measures to get the complete wing area *S* and the full wingspan *b*. Mean chord (m) was then calculated as *c̅* = *S*/*b*, aspect ratio as AR=*b*^2^/*S* and wing loading as WL=*mg*/*S* (N/m^2^), where *g* is gravitational acceleration (9.81 m/s^2^).

#### Statistical analyses

All statistical analyses were carried out using RStudio (4.1.3). First, to test relationship between VeDBA_mean_ and flight duration, as well as between VeDBA_peak_ and flight duration of all flights from all tagged individuals, we used maximum likelihood estimation between the parameters with Weibull distribution, assuming the shape and scale of distribution are influenced by flight duration.

Second, we tested how flight activity, flight performance and wing morphology differed between sexes, which we did using independent samples t-tests.

Wing morphology directly affects flight performance, as the aerodynamic thrust force (**T**) produced by a wing scales linearly with wing area (**T**∼*S*) (Tomotani and Muijres, 2019). Furthermore, Newton’s second law of motion states that a flying bird accelerates proportionally with the thrust force-to-weight ratio (**A**∼**T**/*mg*). Combining these two mechanisms, suggests that the observed VeDBA accelerations scale linearly with the weight-normalized wing surface area, which equals the inverse of wing loading (*S*\*=*S/mg*=1/WL).

We tested this notion using a linaer Pearson Correlation Tests on the relationship between the acceleration metrics (VeDBA_mean,bird_ and VeDBA_peak,bird_) and weight-normalized wing surface area (*S*\*= 1/WL).

In this study, brood size of individual bird was measured on the day of tagging. Foraging activity during chick rearing has previously been shown to correlate significant with brood size (Davies, 1985). Here we tested whether this is also the case for foraging flight behaviour and brood size. For this, we first used two paired sample t-tests on the six breeding pairs, in which we compared the flight proportion and mean flight duration between sexes in each nest box. Second, we used a one-way ANOVA to examine the relationship between clutch size and the flight activity parameters flight proportion and mean flight duration.

Finally, we evaluated weather time of the day influence flight proportions of pied flycatchers. For each individual, one hour after accelerometer device monitoring (i.e., around half of the total accelerometer working time) was selected as the tagging time parameter. We then used a generalized additive model to evaluate relationship between flight proportion and tagging time.

## Results

### Machine learning model for behavioural identification

The classification performance by the XGBoost model can be found in the confusion matrix (Figure 2). Flying and perching has the best overall precisions and recalls. For food consumption related behaviours, swallowing and food shaking were sometimes misclassified between each other. Bill wiping had the lowest recall rate (38.46%) and was often misclassified as food shaking.

**Figure 2.**
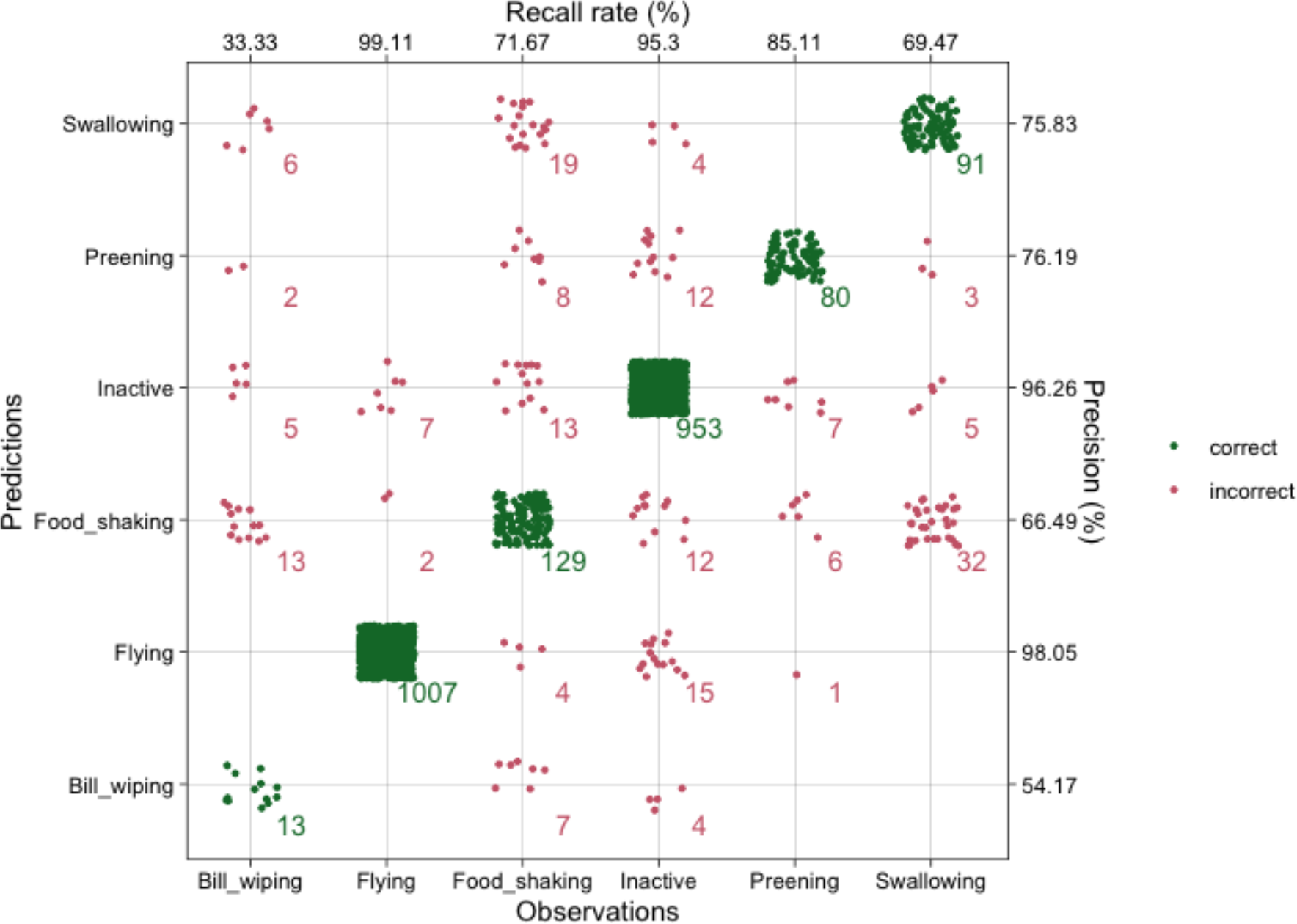
Predic-ons by the XGBoost machine learning model versus ground truth behaviour observa-ons. Here, precision rate = TP/(TP+FP) and recall rate = TP/(TP+FN), where TP, FP and FN are the number of true posi-ves, false posi-ves, and false nega-ves, respec-vely.

### Flight activity, effort, and performance of foraging pied flycatchers

Raw accelerometer data from 26 tagged individuals were classified into six behaviour types by the XGBoost machine learning model, and then converted into flight and non-flight segments (Figure 3). For each flight segment, we estimated the corresponding activity, effort, and performance metrics.

**Figure 3.**
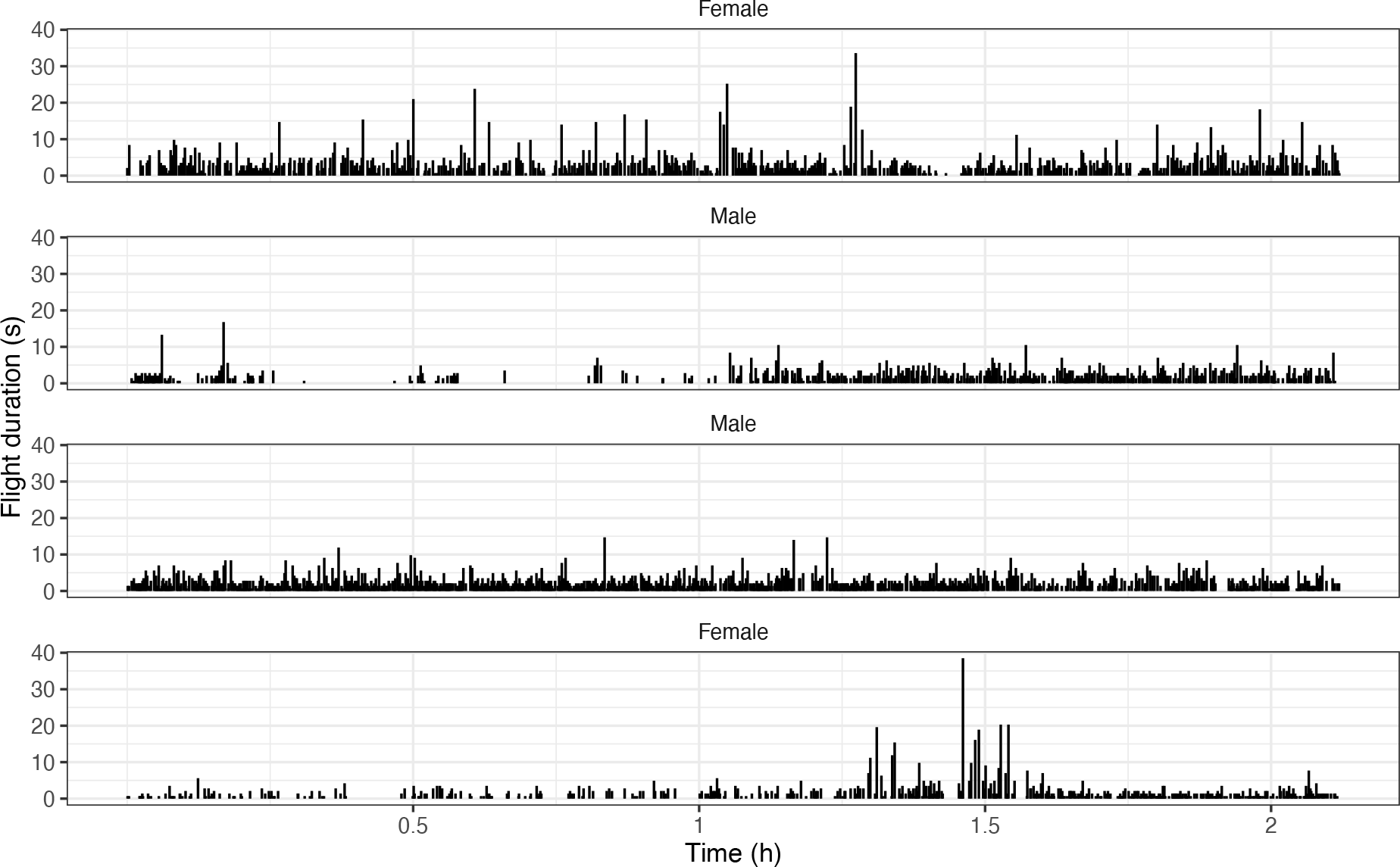
Time allocations and durations of all identified flight segments of four individuals during ∼2 hours accelerometer recordings. Flight activity is detected by the XGBoost machine learning model. Each bar shows the start and duration of a single flight segment (on the abscissa and ordinate, respectively).

### Flight activity: flight proportion and duration

We quantified flight activity using both flight proportion and flight duration (Figure 3). All flights from the posted processed sequences showed large variation of flight proportion and durations between individuals.

The pied flycatchers had a flight proportion that averaged 13.66% of time (n = 26, SD = 6.28%, range = 5.08% - 31.39%). The average number of flights per hour was 199 (n = 26, SD = 92, range = 52 - 549). The mean flight duration averaged 2.38 seconds (n = 26, SD = 0.38, range = 1.64 - 3.15). Short flights with duration less than 10 seconds accounted for 99.29% of all the flights identified from the pied flycatchers among which very short flights that only last for one second was the most frequent flights (Figure 4).

**Figure 4.**
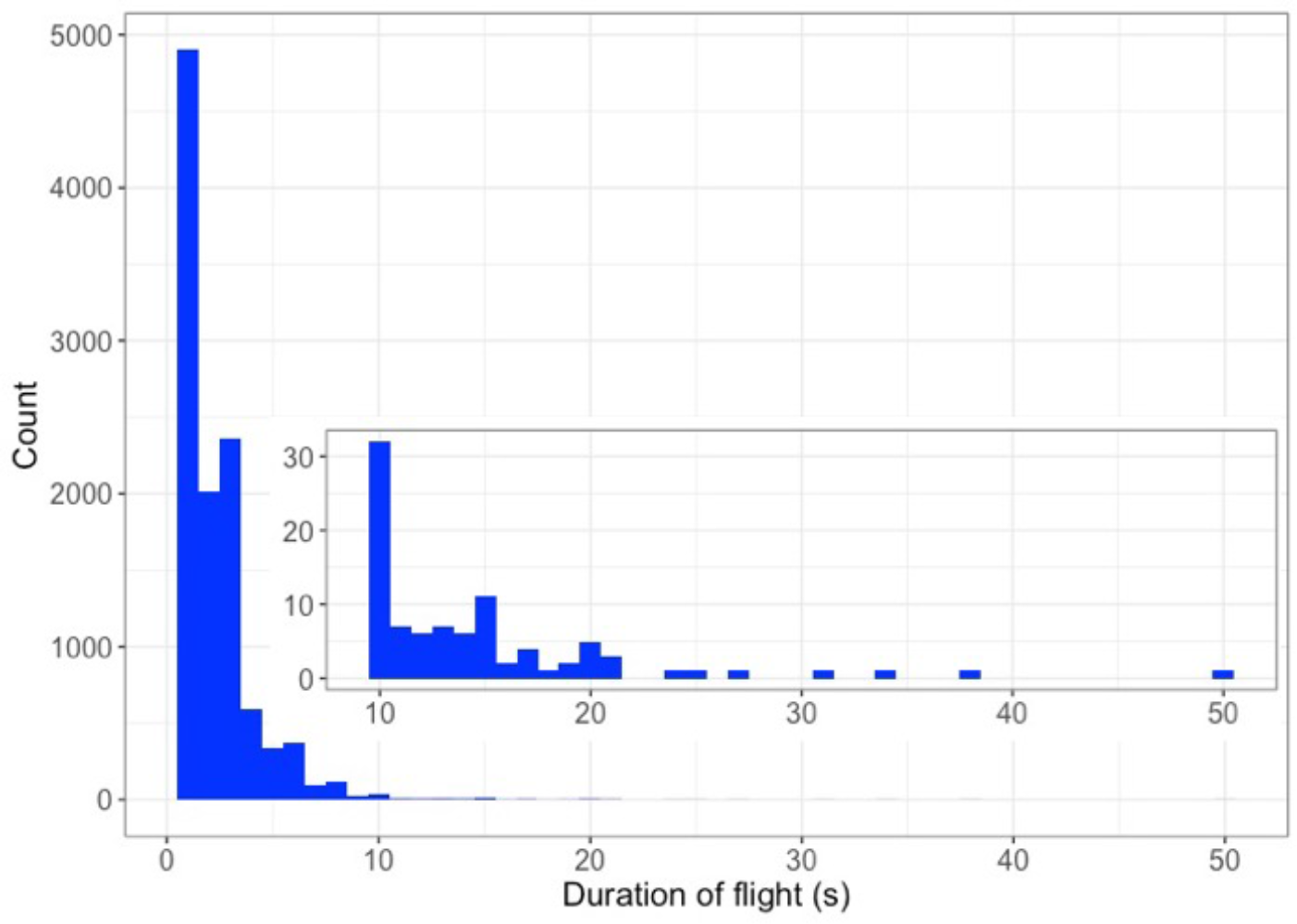
Histogram of the duration of all 10,909 detected flight segments of the 26 foraging pied flycatchers. The inset shows zoomed in data for the flight with a duration of longer than 10 s.

### Flight effort: mean VeDBA during flights

For all the identified flight segments, we estimated the mean acceleration per segment (VeDBA_mean_) as a metric for flight effort (Figure 5A). Short flights had larger variations of VeDBA_mean_, whereas longer flights generally converge to an average VeDBA_mean_ of approximately 0.76*g*, which was supported by the fitted distributions in Figure 5A. Short flights with high VeDBA_mean_ values were most likely flights where the bird flew up with continuous strenuous wing flaps or when performing rapid maneuvers. Slow flights with low mean VeDBAs were possibly associated with descending flights. The long flights with average VeDBA_mean_ were most likely commuting bounding flights at an approximately constant flight speed (Rayner, 1985).

**Figure 5.**
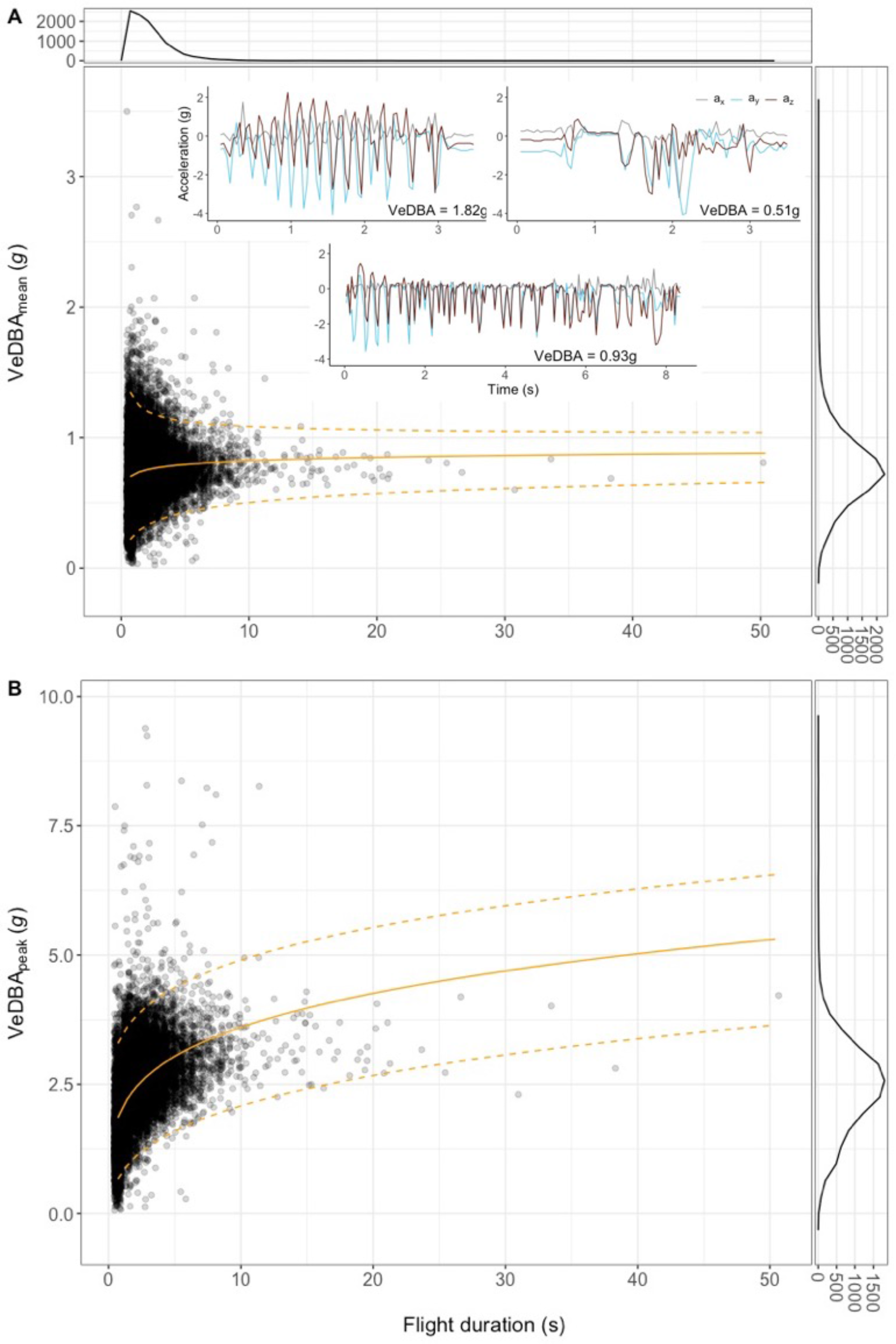
Mean VeDBA VeDBA_mean_(A) and max VeDBA VeDBA_peak_ (B) during all 10,909 flight segments of the 26 pied flycatchers versus flight durations. Each dot represents VeDBA_mean_.(A) and VeDBA_peak_ (B) with the corresponding flight duration of one flight segment (thus, in total 10,909 dots in each panel). The orange lines were fitted by Weibull distributions (i.e., two fits for VeDBA_mean_ and VeDBA_peak_ respectively), with scale and shape parameters influenced by log-transformed flight duration. The solid orange line shows median of fitted distributions, whereas the bottom and top dashed orange lines show 5% and 95% quantiles of fitted distributions respectively. The popup plots in (A) display raw accelerometer data of three flights with high, low and near average VeDBA values. In the low VeDBA panel (i.e., VeDBA = 0.51g), raw accelerometer data of all three axis remain close to 0 *g* around time of 1s. This is due to the wing folding of the bird during flight, which was confirmed in our 2022 aviary experiment. The frequency plot on the top and right show number of occurrence of different flight durations and mean VeDBA (A) and max VeDBA (B) values respectively.

### Flight performance: peak VeDBA during flights

For all the identified flight segments, we estimated the peak acceleration per segment (VeDBA_peak_) as a metric for maximum flight performance during that flight segment (Figure 5B). The distribution of peak total acceleration as expressed by VeDBA_peak_ shows as expected much higher values than for VeDBA_mean_ (Figures 5B and 5A, respectively), with a mean peak acceleration over all flights of 2.44*g* (*n* = 10,909 flights, SD = 0.85*g*, range = [0.06*g* – 9.38*g*]). The number of flights in total that was identified as ‘high-acceleration flight segments’ (VeDBA_peak_>3.5*g*) was on average 36±23 per individual (mean ± standard deviation), with a maximum of 103 high-acceleration segments for one individual. These high-acceleration segments represent on average 9%±5% of the total number of flights (*n*=26 individuals).

### Sex and flight performance

Based on the flight metrics per segment (*n*=10,909 flights), we estimated flight activity, effort, and performance per individual bird (*n*=26 birds), and compared how this differed between sexes (*n*=13 males; *n*=13 females).

### Flight activity: flight proportion and duration

Flight activity did not differ significantly between sexes (Figure 6A-B). Flight proportion (*R*_flight_) did not differ significant between the sexes (Figure 6A; independent t-test on the arcsine square root transformed data: d.f. = 23.38, t = -0.73, p = 0.47). Females flew 13%±7% (*n* = 13) of the time, and males this was 14%±6% (*n* = 13). Also, flight duration did not differ significant between males and females (Figure 6B; independent t-test: d.f. = 23.14, t = -1.00, p = 0.33). For females, flight segments had a duration of *T*_flight_=2.3±0.4 seconds (*n* = 13), and for males this was *T*_flight_= 2.5±0.3 seconds (*n* = 13).

**Figure 6.**
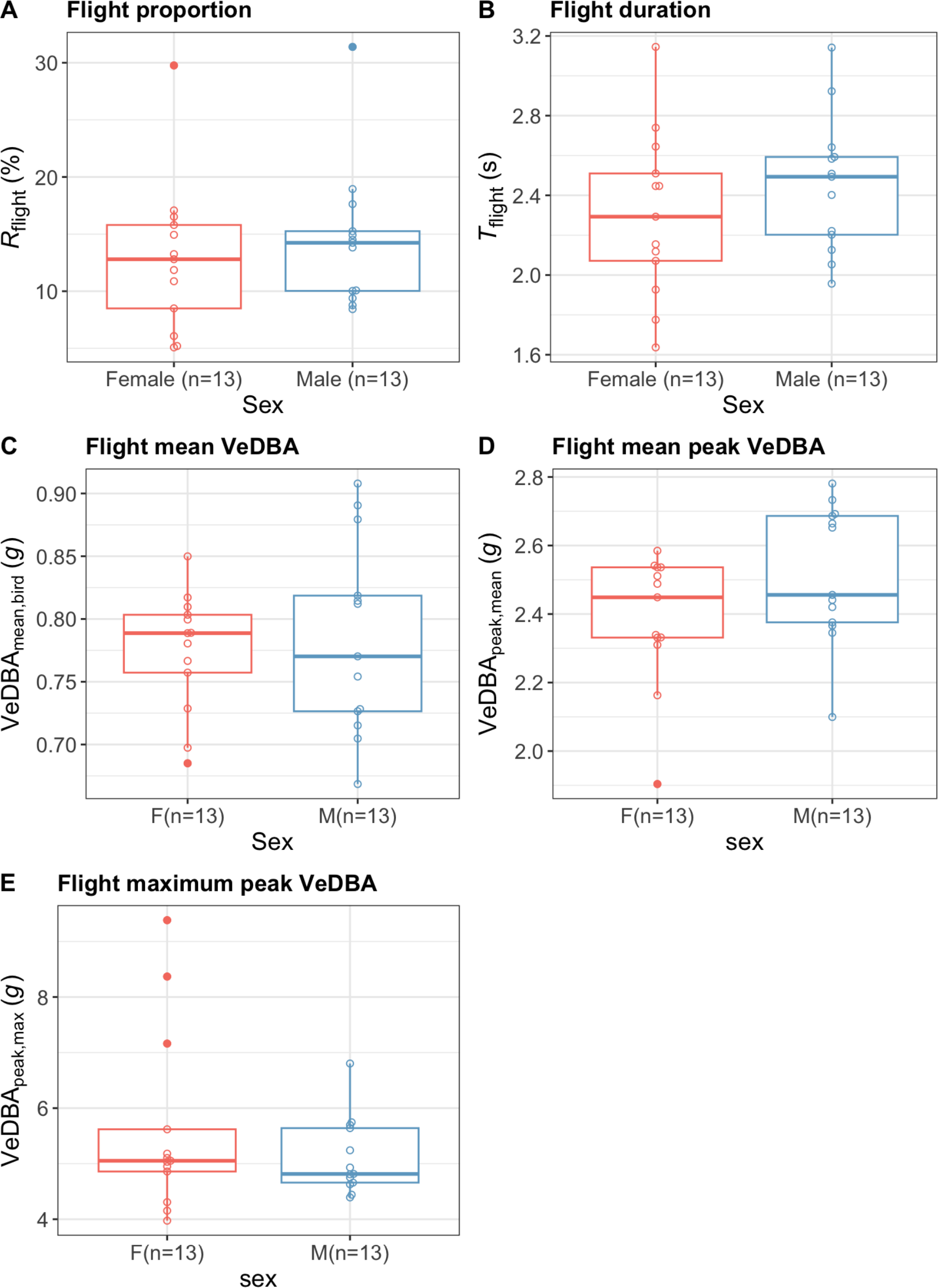
Flight activity, effort, and performance of foraging female and male pied flycatchers, estimated using accelerometer recordings in the field. All data for females (F) and males (M) are in red and blue, respectively. The flight parameters include (A) flight proportions *R*_flight_, (B) mean flight duration per flight segment *T*_flight_, (C) mean in-flight accelerations per bird VeDBA_mean,bird_, (D) mean peak in-flight accelerations per bird VeDBA_peak,mean_, (E) and the maximum peak in-flight accelerations per individual VeDBA_peak,max_.

Finally, we performed paired t-tests on the flight activity metrics per pair (nest box), to test whether flight activity of the male and female in a single pair depends on each other. For this, also no significant difference was found between sexes in either flight proportion (arcsine transformed paired t-test: d.f. = 5, t = -0.83, p = 0.44) or mean flight duration (paired t-test: d.f. = 5, t = -2.2, p = 0.08).

### Flight effort: mean in-flight VeDBA per bird

Flight effort as quantified using mean VeDBA did not differ significantly between males and females (Figure 6C; independent t-test: d.f. = 20.02, t = 0.36, p = 0.72). Females had an in-flight mean VeDBA of 0.77±0.05*g* (*n* = 13), and for males this was VeDBA_mean,bird_=0.78

±0.08*g* (*n* = 13).

### Flight performance: peak in-flight VeDBA per bird

Flight performance also did not differ significantly between males and females (Figure 6D- E). This was the case for both the mean and maximum peak VeDBA (Figure 6D and 6E, respectively; independent t-test between sexes for VeDBA_peak,mean_: d.f. = 23.94, t = -1.69, p = 0.10; independent t-test between sexes for VeDBA_peak,max_: d.f. = 16.01, t = 1.02, p = 0.32).

During their foraging flights, females had a mean peak VeDBA of 2.39±0.19*g* (*n* = 13) and for males VeDBA_peak,mean_=2.52±0.20*g* (*n* = 13). The corresponding maximum peak VeDBA per individual female was 5.63±1.65*g* (*n* = 13), and for individual males their maximum peak VeDBAs were 5.12±0.69*g* (*n* = 13).

### Sex and morphology

None of the primary body and wing morphology parameters differed significant between the males and females (Fig. 7A-D). Body mass did not differ between males and females (males: *m*=13.0±0.49 g; females: *m*=13.2±0.63 g; independent t-test, df = 21.86, t = 0.84, p-value = 0.41; Figure 7A). Also, females and males had non-significant different aspect ratios of AR=4.98±0.38 and AR=5.14±0.3, respectively (independent t-test, df = 15.05, t = -1.07, p = 0.3; Figure 7B).

**Figure 7.**
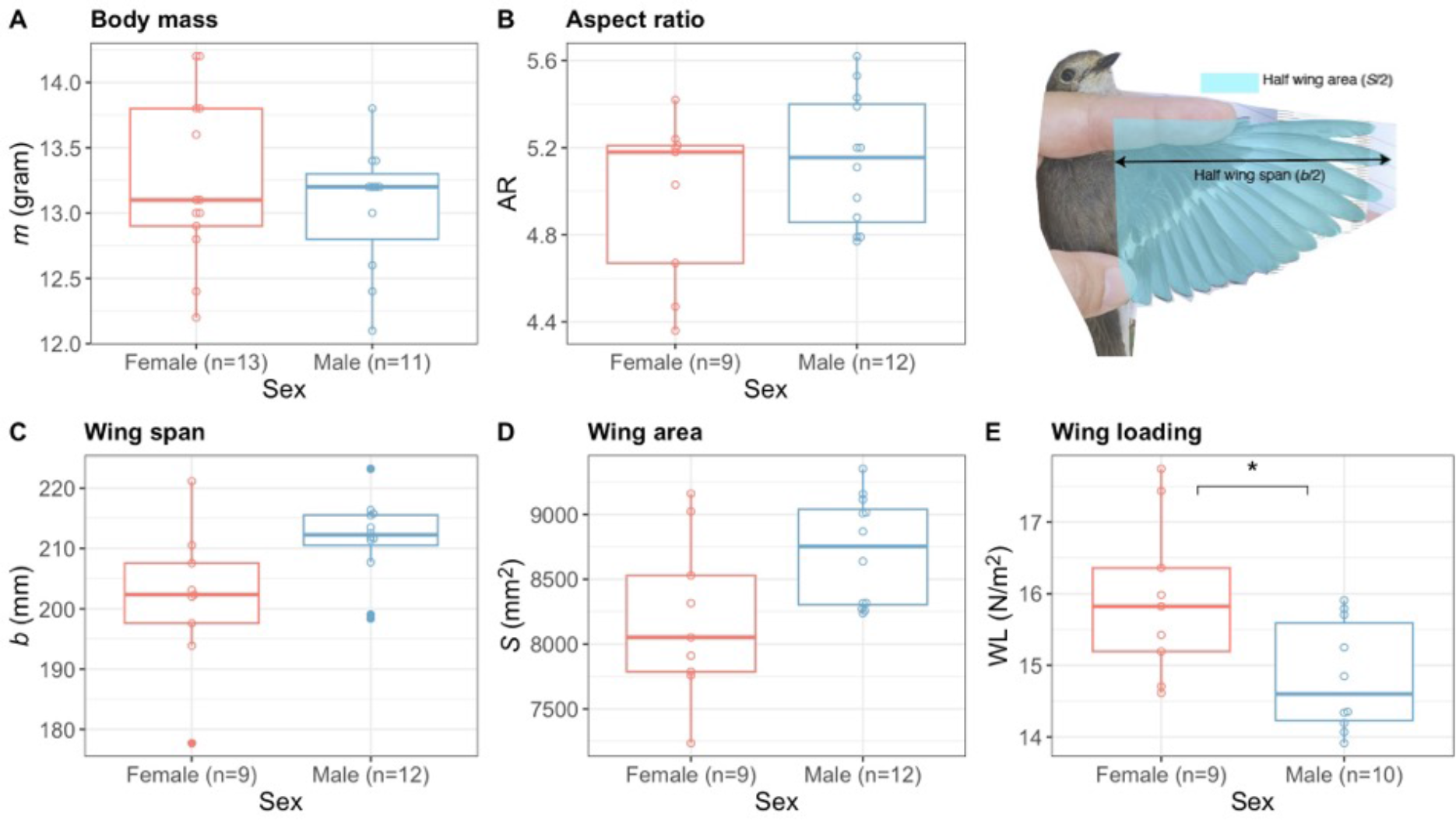
Morphological differences between the studied female and male pied flycatchers. All data for females (F) and males (M) are in red and blue, respectively. Measurements include (A) Body mass *m*, (B) wing aspect ratio AR, (C) wing span *b*, (D) wing area *S*, (E) Wing loading WL. We estimated semi-span (*b*/2) and half the wing surface area (*S*/2) based on digital images of the bird wing (top right).

Although also not significant, the males appeared to have larger wings (longer wingspan and larger wing area) than females. Males appear to have a ∼4% larger wing span (males: *b*=211±7 mm; females: *b*=202±12 mm; independent t-test, df = 12.07, t = -2.16, p = 0.052; Figure 7C), and a ∼6% larger wing area (males: *S*=8712±418 mm^2^; females: *S*=8196±625 mm^2^; independent t-test, df = 13.19, t = -2.14, p-value=0.051; Figure 7D).

Despite these non-significant differences in the primary body and wing morphology parameters, the derived wing-loading parameter did differ significant between males and females (Figure 7E). Females had on average a ∼7% larger wing loading than males (females: WL=15.9±1.11 N/m^2^, males: WL=14.8±0.77 N/m^2^; independent t-test, df = 14.11, t = 2.45, p-value = 0.028).

### Morphology and flight performance

Wing loading directly affects the ability to produce in-flight accelerations, and thus we focussed primarily on how wing loading affects flight activity, effort and performance. We did not find any significant relationship between wing loading and the flight-activity metrics flight proportion and flight duration (Pearson correlation test on WL versus *R*_flight_: df = 17, t = -0.008, p =0.99; Pearson correlation test on WL versus *T*_flight_: df = 17, t = 0.33, p = 0.75).

Because in-flight accelerations scale inversely with wing loading, we evaluated the relationship between wing loading and the VeDBA-based metrics using a linear regression on the inverse of wing loading, called weight-normalized wing area (*S*\*=1/WL, Fig. 8). This shows that both mean and peak VeDBA correlate significantly and positively with weight-normalized wing area (Pearson correlation test on *S*\* versus VeDBA_mean,bird_: p = 0.011, *R*^2^ = 0.32; Pearson correlation test on *S*\* versus VeDBA_peak,mean_: p = 0.003, *R*^2^ = 0.41).

**Figure 8.**
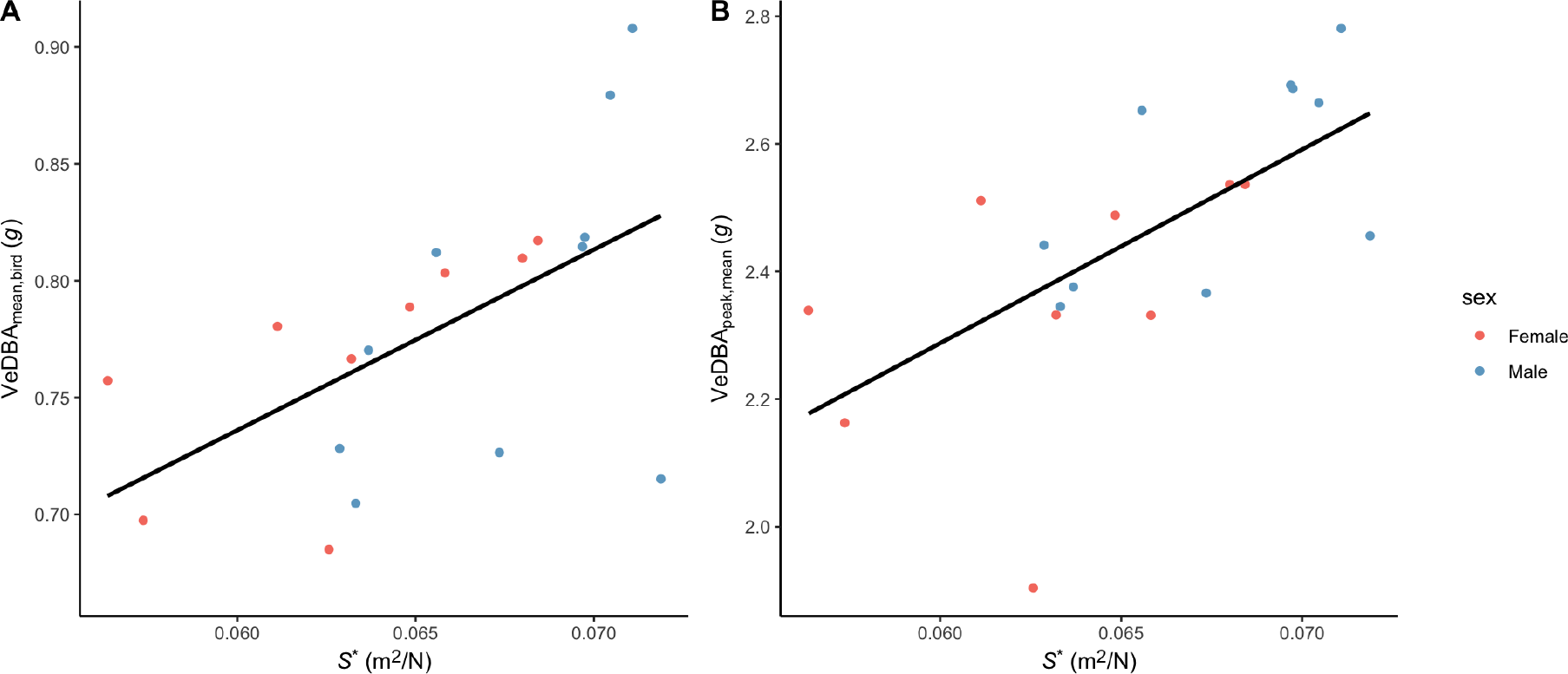
In-flight acceleration versus the weight-normalized wing area *S*\* for all studied birds. All data for females (F) and males (M) are in red and blue, respectively. In-flight accelerations are quantified using (A) mean accelerations per individual (VeDBA_mean,bird_), and (B) the corresponding average peak values per individual (VeDBA_peak,mean_). Data points show the values per individual, color-coded with sex. Black lines show the significant linear regression between the acceleration metric and weight-normalized wing area.

Thus, among all tested birds both flight effort and flight performance (as expressed by VeDBA_mean,bird_ and VeDBA_peak,mean_, respectively) scale linearly and positively with weight- normalized wing area, and thus inversely with wing loading. Because males have on average lower wing loading than females (Fig. 7E), they should also be able to produce high in-flight accelerations, but they did not during the sampled foraging periods (Fig. 6C-E).

### Brood size and flight performance

We tested for a possible relationship between brood size (range from 3 to 8 chicks) and flight activity. The flight activity metrics flight proportion and flight duration per flight segment did not depend on brood size (One-way ANOVA on arcsine transformed *R*_flight_; F-value = 0.72, p-value = 0.62, Figure 9A; One-way ANOVA on *T*_flight_: F-value = 1.24, p-value = 0.33, Figure 9B).

**Figure 9.**
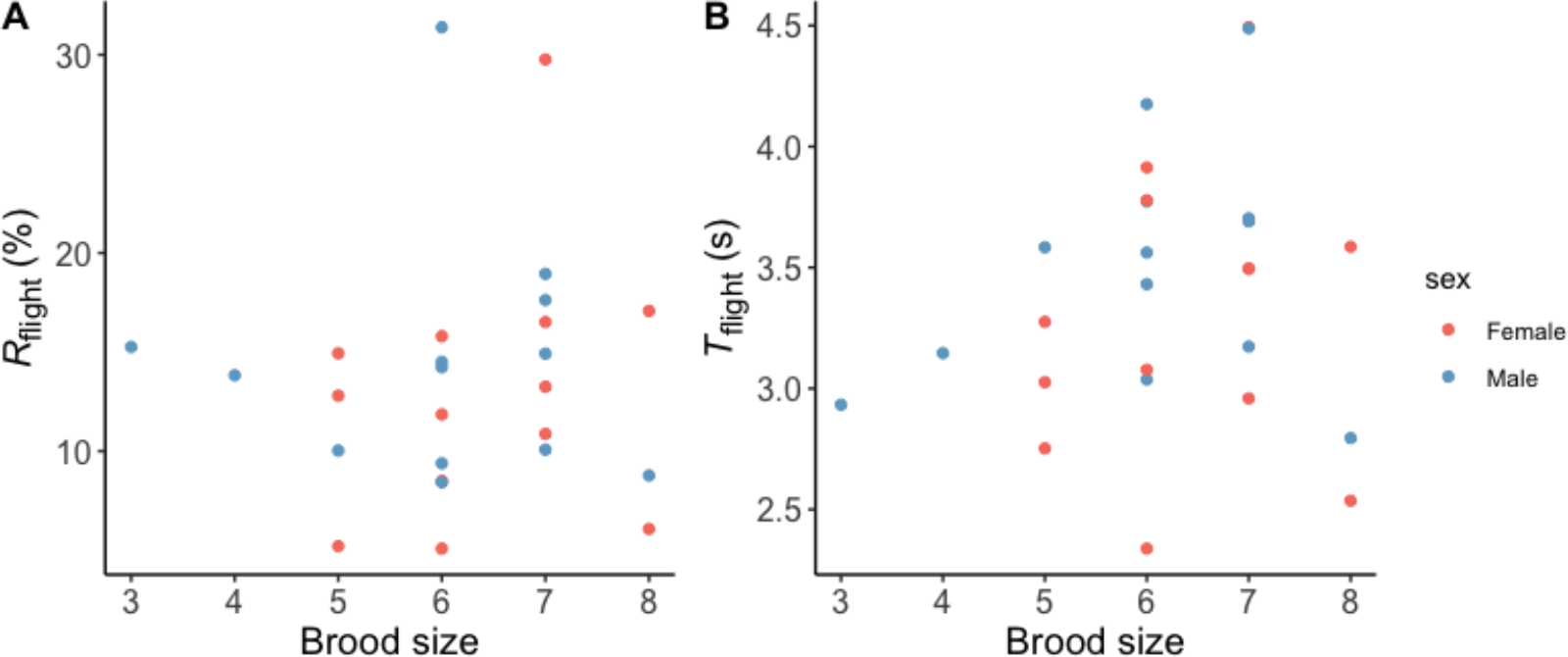
Flight activity versus the brood size as number of nestlings for all studied birds. Flight activity was quantified using (A) flight proportion per individual (*R*_flight_), and (B) the mean flight duration of all flight segments per individual (*T*_flight_). Data points show the values per individual, color-coded in red and blue for females and males, respectively.

### Time of day and flight performance

Finally, we tested how flight activity varied during a day of foraging, by correlating the average flight proportion per individual with time of day (Fig. 10). From the gam test, relationship between flight proportion and time of day was not significant (F = 2.08, p = 0.16). However, the fit result shows that foraging flight activity is highest in the morning, when birds flew approximately 15% of the time. Flight activity dips in early afternoon to 8%, and then increases again towards the late afternoon.

**Figure 10.**
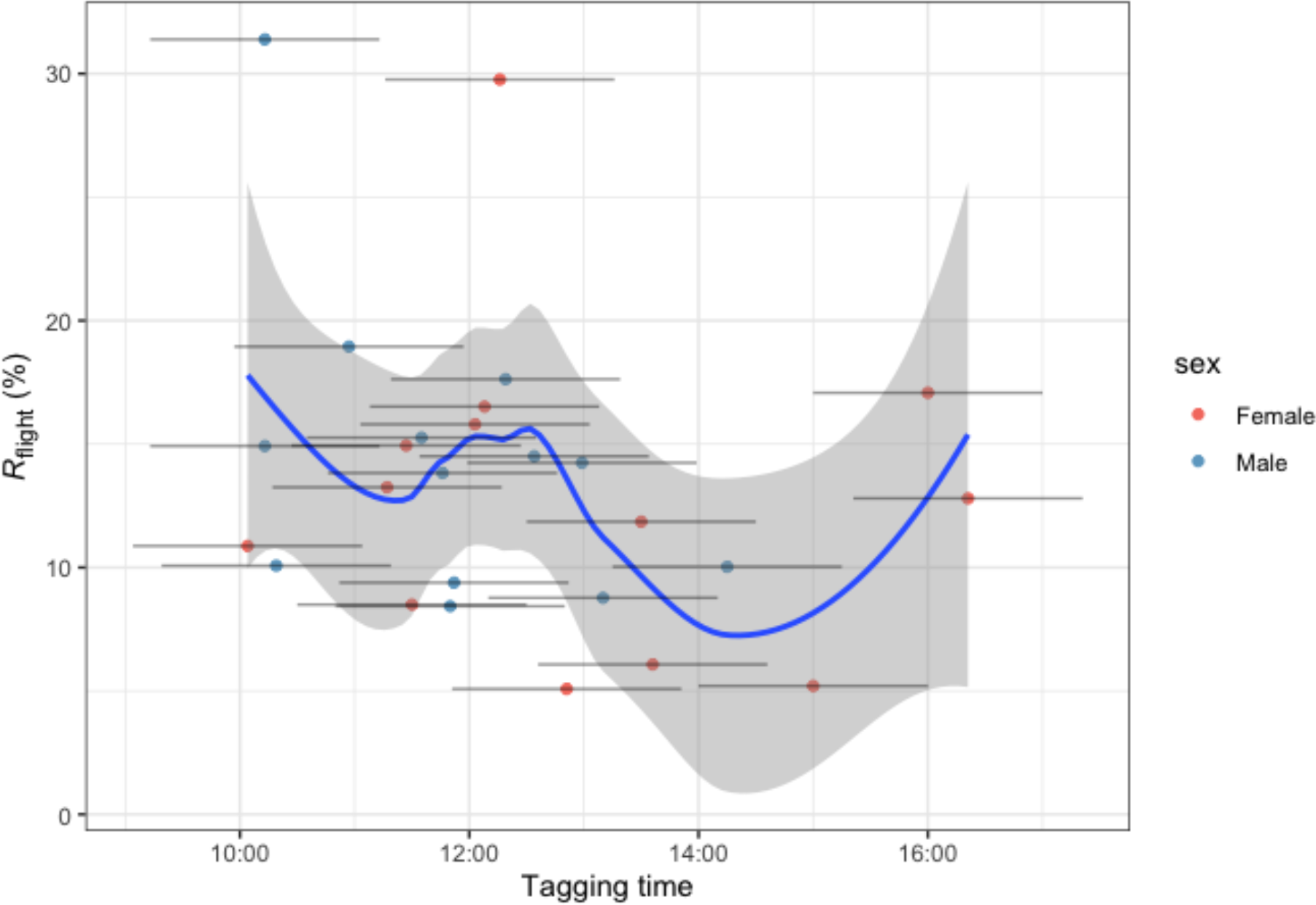
Flight proportion (*R*_flight_) in relation to time of day of the sampling (tagging time). The horizontal bars show the sampling period for each individual; the corresponding data points show the mean tagging time per sampling used for the curve fitting; data points are color-coded in red and blue for females and males, respectively. The blue curve shows the loess fit on the data, and the grey shading represent the confidence interval of the curve.

## Discussion

### Foraging flight activity, effort and performance

#### Flight activity

The pied flycatchers in our study were actively flying on average 14% of the time during the two-hour window of our observations. At first this may seem low if considering that these birds are regarded as very active flyers. The birds spent their time rather than flying and being inactive into preening, feeding, and possibly feeding chicks. It is worth noting that the tagged birds might still be recovering from the disturbance of being captured, handled and tagged.

In a theoretical paper on optimal fuel loads and stopover use during migration a general proportion of flight and stopover during the complete migration was estimated to be 1:7 (Hedenström and Alerstam, 1997), which arises simply as the ratio between power required to fly and the rate of energy accumulation. Interestingly, our flycatchers exhibit very similar proportion, 1:7.3. It may of course be a coincidence because the condition the birds live under during breeding is vastly different from when on migration, but it is also possible that this stems from a general ratio between time required to feed fuel to cover a unit flight time, no matter if it is migratory or transport flight.

#### Flight durations and number of flights

Flight durations and number of flights of breeding songbirds have rarely been studied previously. Among the tagged pied flycatchers, although there were occasional flights of 30- 50 seconds, most flights were below 10 seconds in length. The average number of flights per hour was 199. This gives us a picture of how these birds are moving through the habitat, performing many short flights. With so many flights it is not likely that the birds only use a sit-and wait foraging strategy typically associated with these birds. With on average more than three flights per minute, the flycatchers probably used short flights to move through the territory searching for food and only occasionally make aerial forays to catch prey. Indeed, previous studies have shown that prey types that pied flycatcher parents provide to their nestlings include Lepidoptera, Coleoptera, Arachnida, Diptera, Hymenoptera, Odonata and Isopoda, among which larvae of the insects that could not fly seemed to be the dominant food type for nestlings (Lundberg and Alatalo, 2010). The ‘high-acceleration flight events’ that we defined as having peak VeDBA of larger than 3.5*g*, most likely represent aerial prey capture events. Among our flycatchers, 9% of all flights consisted of these high-acceleration flights, which includes per individual 36 high-acceleration flights during the two-hour recording period.

The many short flights have a potential implication for the energy expenditure of the flycatchers. Every time the bird takes off and lands, it first has to accelerate from a stationary position, which takes a lot of energy (compared to steady flight) (Nudds and Bryant, 2000). During the consecutive landing manoeuvre, the animal needs to produce large wing forces to brake before touch-down, as well as buffer the forces with the legs and feet at touch-down (e.g., Provini et al., 2014). Changing speed like this frequently must come with added energy expenditure as well as requiring skilful manoeuvring while flying in a complex and dense forest environment (cf. Henningsson, 2021).

Among all flights, short flights (i.e., <10 s) had larger variations of flight effort, as expressed by mean VeDBA. VeDBA values were found positively related to energy expenditure of flying birds (e.g., Sutton et al., 2023). Therefore, the flights with large VeDBA values possibly associated with ascending flights or rapid flight manoeuvres, which both require increased energy expenditure. On the other hand, there were also large number of short flights with low VeDBA values, possibly associated with descending flights, or short flights between tree branches. For flights longer than 10 s, mean VeDBA values were close to mean VeDBA values of all flights, indicating energy conserving strategy of bounding flights at constant speeds. Our raw accelerometer data confirm the use of such bounding flights during these long flight segments, as the alternating active flapping and wing folding phases can be observed.

#### Total flight distance per day

Based on flight proportion, we can estimate the total fight duration and distance travelled by the birds during a day of foraging. As the birds flew on average 13% of the two hours of monitoring, they were in flight for approximately 15 minutes. We can extrapolate from this to get a rough estimate of the total flight duration and distance in a single day.

The foraging hours of breeding pied flycatchers have been shown previously to be approximately 17 hours per day, from approximately 4:00h in the morning to 21:00h in the evening (Lundberg and Alatalo, 2010). During this period, the feeding rate was strikingly constant. With a 13% flight percentage, our pied flycatchers were thus actively flying for approximately 2 hours and 13 minutes during the daily 17 hours of foraging.

The average flight speed of migrating pied flycatchers have been measured previously to be 9.74 m/s (Alerstam et al., 2007). This estimate was during migration, and the flight speed during foraging and commuting in its home range might in some cases be lower and sometimes higher. But if we assume this migration speed to be roughly equal to the average speed during foraging, then we can estimate the total flight distance during a single day per individual, based on the two-hour flight activity of that bird. For the 26 monitored birds, the total distance travelled during a day of foraging was 84±38 km, ranging from 31 to 192 km. According to the Swedish ringing recovery data summarized by Ellegren (1993), pied flycatchers have an average migration speed of approximately 60 km/day in autumn (n = 19), and the highest migration speed reached 116 km/day. Thus, this back-of-the-envelope calculation tells us that the flycatcher’s flight distance in one day during breeding is similar to the distance travelled per night of migration. We often view the migration of the long-distance migrants as the most impressive feat they do during the annual cycle, but the feeding of the young should not be overlooked in terms of flight performance.

#### Flight effort and performance based on in-flight accelerations

The mean peak VeDBA among all individuals was 2.5*g*. This can thus probably be assumed to represent the magnitude of the *g*-force that the bird is exposed to during flapping motion in flight. Similar accelerations in straight and level flight have been reported for a barn swallow flying in a wind tunnel (Pennycuick et al., 2000), so this probably is not extreme but simply a consequence of the flapping wing propulsion.

If we instead look at the highest accelerations that the birds generated, we find that the recorded maximum peak VeDBA was 9.4*g*. If we put this in perspective, 9*g* is what modern jet fighters are designed to handle at most. So, when the flycatchers are pushing their performance like this, presumably during aerial prey catching events, they are performing rather dramatic manoeuvres that put a lot of strain on their bodies. However, one should note that these events are brief, which is probably why it is possible for the birds to manage it.

This is confirmed if we look at these ‘high acceleration flight events’ in our dataset, in which the number of consecutive flight seconds with mean peak accelerations above 4.5g, across all individuals, was on average 2.5 seconds. On the other hand, the total number of high acceleration flights was on average 37 during the two-hour period, with a maximum of 149, so the birds are frequently exposed to these short high accelerations in their normal routine behaviour.

### Morphology and flight performance

Males had longer wingspan and larger wing area than females, which is consistent with a previous study showing that males have longer wingspan than females (de la Hera et al., 2014). While the aspect ratio between the two sexes was similar, males had significantly lower wing loading. Wing loading directly affects the accelerations that a flying bird can produce, as aerodynamic thrust forces scale with wing area, and in-flight accelerations scale with the ratio between thrust and body mass (Pennycuick, 1989). Thus, flight performance as quantified by in-flight accelerations (VeDBA) should scale inversely with wing loading (WL) as VeDBA∼1/WL. Our data of 26 foraging pied flyctachers confirm that this is also the case for both mean and peak accelerations during foraging flights in the field. It is surprising though, that despite their lower wing loading, male flycatchers do not exhibit higher VeDBA values during foraging and chick rearing than the female birds. This suggest that although male pied flycatchers are capable of producing higher in-flight accelerations than females, they do not do so and thus operate at a reduced effort.

We can model the relative effort exhibited by the males and females using the linear regression between in-flight accelerations and the weight-normalized wing surface area (Fig. 8A). This regression shows that VeDBA scales linearly with the inverse of wing loading as

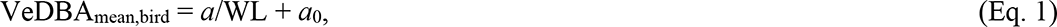

where *a* and *a*_0_ are the slope and intercept of the linear regression, respectively (*a*=7.73 and *a*_0_=0.27, figure 8A). The model is in line with the functional relationship between wing loading and in-flight accelerations, according to aerodynamic theory (**T**∼*S*) and Newton’s second law of motion (**A**=**T**/*m*), leading to **A**∼1/WL. Applying this model to the mean wing loading of male and female pied flycatchers, allows us to estimate the average VeDBA_mean,bird_ that the male and female flycatchers can produce during foraging flights. Comparing these values provides an estimate of their relative in-flight foraging effort.

Males have on average a 7% lower wing loading than females (males: WL=14.8; females: WL=15.9; Fig. 7E), which enables them to produce 5% higher mean in-flight accelerations (Eq. 1; males: VeDBA_mean,bird_ = 0.792*g*; females: VeDBA_mean,bird_ = 0.756*g*). Thus, for males and females to produce similar in-flight accelerations during foraging flights for chick rearing, the female flycatchers need to effectively operate at a 5% increased relative effort compared to their male partner.

### Foraging behaviour and territory size

We found that 99% of flights had a duration below 10 s. If we use the previous estimate of flight speed by Alerstam et al. (2007) of 9.74m/s and make the assumption that the birds fly out from the nest and back again in two main flights that means that 99% of the flights were within 100 m radius of the nest. If we use the mean flight duration of 2.38s it means that the average radius is around 23m. Of course, it could also be the case that the birds fly several consecutive 100-meter flights away from the nest and back again and then may reach beyond this radius. So one should keep in mind that this is a rough estimate and merely an indication of habitat use. However, if we accept the assumption that it is more likely that the birds only make single flights since the availability of insect food can be assumed to be high and easy to find, this information may still be important in telling us the movement of the birds in their habitat and grants us the possibility to get a better picture of the size of the core home range for these birds. From the perspective of conservation biology, this indicates how large territories need to be to sustain a breeding pair. Obviously, it will vary with the quality of the territory (Bibby and Green, 1980), but it gives a rough idea about area size needed. How does this compare with other species that feed on different types of food – for example seed eaters? This needs to be addressed in future studies and would grant invaluable information about breeding bird populations to be used when planning nature reserves or other types of protection of habitats. On the other hand, 19 out of 26 individuals took occasional longer flights over 10 s and 5 individuals had flights longer than 20 s. These longer flights might be explorational for better foraging habitat or in males seeking for unoccupied nest box for pairing with a secondary female (Alatalo et al., 1982, Lamers et al., 2020).

### Breeding performance and flight activity

No significant difference was found either for flight proportion or mean flight duration of parent birds in relation to brood size. This could partly be due to the rather low spread of the clutch sizes, including only clutch sizes from three to eight, and with most samples within five to seven chicks. Future studies could test this by experimentally manipulating brood size. Nevertheless, if for argument’s sake, no difference was genuinely the case, we could speculate that in natural broods the parents may not work as hard as they can in order to preserve energy (due to parent-offspring conflict), which is similar to the case studied by Masman et al. (1989) in kestrels. They found in their study that male kestrels spent similar proportion of time in flight irrespective of brood size, but their flight-hunting yield (prey captured per hour flight-hunting) was positively correlated with brood size, to guarantee their nestlings had enough food (Masman et al., 1989). However, when kestrels met a situation of food shortage, males allocated more time to flight (Masman et al., 1989). In our case, flycatcher parents with larger brood sizes might either work more efficiently to bring more food back to chicks in each visit (Siikamäki et al., 1998), or they occupy better territories where the same flight effort results in more prey.

### Division of parental care labour between sexes

The paired sample t-tests that we did including six nest boxes showed no significant difference in either flight proportion or mean flight duration between sexes within pairs. This result supports the conclusion that there is no difference in parental investment between the sexes, if it comes to time and activity investment. These tests are relevant since by comparing between male and female of the same nest, we can be sure that they have the same ‘task at hand’ since they are feeding the same chicks and therefore face the same challenge. However, in some cases the male may leave the nest and his primary female to occupy a new territory and try to mate with a secondary female (Lundberg and Alatalo, 2010). In polygynous pairs, the parental care the male provides might be less than that of the female, especially in secondary nests (Alatalo and Lundberg, 1984). There were only two polygynous cases in our dataset, and only adult females (secondary females of polygamously mated males) in these two broods were caught and deployed with accelerometers. The frequency of polygyny varies a lot in pied flycatchers between different populations (Lundberg and Alatalo, 2010). More data should be collected in future studies from pairs of broods where polygyny occurs if we would like to test the flight proportion (and mean flight duration) difference between polygynous and monogamous males.

Our data thus showed that female and male partners invest a similar amount of flight time into raising offspring, but our study also allowed us to estimate how much effort the male and female flycatchers invested during these foraging flight periods. Both in-flight acceleration metrics VeDBA_mean_ and VeDBA_peak_ did not differ between males and females. As these parameters were used as metrics for quantifying flight effort and performance, respectively, this might suggest that also flight effort did not differ between sexes. But as mentioned above, male pied flycatchers have a significantly lower wing loading than females, which should allow them to produce higher VeDBA_mean_ and VeDBA_peak_ values during in-flight foraging. The fact that they do not shows that male pied flycatchers invest relatively less effort in chick rearing than females, as they operate at lower accelerations compared to their potential.

Several previous studies have suggested that male and female pied flycatchers invest similar effort in chick rearing (Alatalo et al., 1988, Siikamäki et al., 1998). By including a validated model for quantifying the effect of wing loading on flight effort in our analysis (Eq. 1), we show that this apparent equal contribution of males and females to parental care is the result of unequal investments in effort. Due to their lower wing loading and consequent higher baseline flight performance, male pied flycatchers might achieve the same parental care output as females, but at an approximate 5% lower effort compared to the females. This shows that the balance in parental care between male and female songbirds is more intricate and complex than previously thought.

## Acknowledgements

We are grateful to Arne Andersson and Johan Bäckman at Department of Biology, Lund University for developing the micro data loggers used in this study and for setting them up before measurements and downloading data from them after.

## Author contributions

Conceptualization: H.Y, F.T.M., A.H., P.H.; Methodology: H.Y., S.L., F.T.M., P.H.; Formal analysis: H.Y., S.L., H.J.K., P.H.; Investigation: H.Y., S.L., J.S.L., A.H., K.P.L, P.H.; Resources: H.Y., K.P.L., P.H.; Writing – original draft: H.Y., S.L., P.H.; Writing – review & editing: H.Y., F.T.M., J.S.L., H.J.K., A.H., K.P.L., P.H.; Visualization: H.Y.; Supervision: F.T.M., H.J.K., A.H., P.H.; Project administration: A.H., P.H.; Funding acquisition: F.T.M., H.J.K., A.H., P.H.

## Data availability

Data used for analyses will be deposited at Figshare upon manuscript acceptance.

